# A Neural Mechanism for the Opportunity Cost of Time

**DOI:** 10.1101/173443

**Authors:** Sara M. Constantino, Jessica Dalrymple, Rebecca W. Gilbert, Sara Varanese, Alessandro Di Rocco, Nathaniel D. Daw

## Abstract

Recent interest has focused on a class of decision problems in which subjects encounter options serially and must decide when to leave an option in search of a better one, rather than directly comparing simultaneously presented options. Although such problems have a rich history in animal foraging and economics, relatively little is known about their neural substrates. Suggestively, however, a separate literature has argued that the key decision variable in these tasks – the opportunity cost of time, given by the average reward rate – may also govern behavioral vigor and may be reported by tonic dopamine (DA).

In this study, we test whether this putative dopaminergic opportunity cost signal plays an analogous role in serial decisions by examining the behavior of patients with Parkinson’s disease (PD), on and off their DA replacement medication, in a patch-foraging task. In these tasks, subjects’ decisions about when to leave a depleting resource implicitly reflect their beliefs about the opportunity cost of time spent harvesting that resource. Consistent with the opportunity cost hypothesis, umedicated patients harvested longer than matched controls, and medication remediated this deficit. These effects were not explained by motor perseveration. Our results suggest a functional role for DA, and an associated cognitive deficit in PD, in a type of decision process that may be distinct from (but related to) the neuromodulator’s well studied roles in behavioral invigoration and learning from rewards.

**Significance Statement:** This study addresses two important questions whose answers are, unexpectedly, linked. First, what is the scope of cognitive functions of the neuromodulator dopamine, whose contributions – for instance, as assessed by both the motoric and more subtle cognitive deficits of patients with PD, which depletes dopamine – range from movement to reward and decision-making? Second, what are the neural mechanisms supporting an important but understudied class of problems, in which, rather than choose among a set of alternatives (like apples and oranges), one makes serial decisions about whether to stick with an option (like a job, or a mate) or seek another? We demonstrate a novel cognitive deficit in PD that integrates this function into the web of DA’s contributions.

## Introduction

Decision neuroscience research has focused on the choice between simultaneously presented alternatives (Ito and Doya, 2009; Barraclough et al., 2004; Tom et al., 2007; Krajbich et al., 2010). However, there has been recent interest in decisions where options are encountered serially and the choice is whether to engage with the current prospect or seek a new one. Such decisions arise in search problems (for jobs, mates, or internet results) and feature in the classic animal foraging literature (Pyke, 1980; Kacelnik, 1984; Stephens and Krebs, 1986). The central idea in foraging is that serial decisions are about time allocation: Engaging with an option is worthwhile only as long as received rewards exceed what you would otherwise expect to earn during that time. In this class of problems, this expectation — the <u>opportunity cost of time</u> — optimally equals the long-run average reward rate in the environment (Charnov, 1976; Stephens and Krebs, 1986).

Relatively little is known about the neural basis of serial stay-switch decisions, although researchers have recently argued that they engage distinct cortical mechanisms from extensively studied simultaneous choice (Hayden et al., 2011; Kolling et al., 2012). A separate, related literature has centered on decisions about effort: How vigorously to work (Robbins, 1976; Niv et al., 2007; Guitart-Masip et al., 2011; Niyogi et al., 2014). Such decisions should in theory be governed by parallel opportunity cost considerations, since sloth becomes more costly as earnings potential increases. Niv et al. (2007) suggested that tonic DA might signal the average reward rate and directly modulate vigor. This idea accords with (and reconciles) the involvement of DA in signaling rewards *and* invigorating movement; It extends the prominent hypothesis that <u>phasic</u> DA signals reward prediction errors for learning (Schultz et al., 1997) by noting that the average of the prediction errors, possibly reflected in accumulated tonic extracellular DA, is equal to the average reward. The proposal that DA mediates the coupling between experienced reward rate and behavioral vigor has been supported by pharmacological studies investigating reaction times (Beierholm et al., 2013) and reach speeds (Mazzoni et al., 2007).

Since foraging and vigor share their central decision variable, we hypothesized that this putative dopaminergic opportunity cost signal might play an analogous role in foraging decisions — in this case, governing the reward level at which people leave a prospect in search of another. To test this, we studied the foraging decisions of healthy volunteers and patients with PD (Dauer and Przedborski, 2003), both on and off dopaminergic medication. Several studies have shown reinforcement learning deficits in PD during simultaneous choice (Frank et al., 2004; Shohamy et al., 2004; Frank, 2005; Cools et al., 2007), and have interpreted them in terms of phasic reward prediction errors, but none has examined the hypothesized role of tonic DA in serial switching.

In this study, subjects repeatedly harvested apples from trees with diminishing returns and had to decide when to leave a tree for a replenished one. In such tasks, animals and people reliably adjust the level at which they leave an option according to the average reward rate, which is consistent with the optimal model (Cowie, 1977; Pyke, 1980; Kacelnik, 1984; Smith and Winterhalder, 1992; Jacobs and Hackenberg, 1996; Thompson and Fedak, 2001; Hutchinson et al., 2008; McNickle and Cahill, 2009; Constantino and Daw, 2015; Wolfe, 2012). We predicted that if the average reward rate is signaled by tonic DA, then PD patients (off dopaminergic medication) would stay with trees longer than controls, reflecting a lower implicit opportunity cost of time, and that medication would ameliorate this deficit.

Our findings suggest a role for DA in controlling behavioral switching, contributing to the emerging literature on the neural mechanisms of foraging and serial decision-making, and reveal an associated cognitive deficit in PD. Moreover, via the notion of opportunity cost, they highlight the conceptual relationship between this cognitive effect on decisions and DA’s classic involvement in movement invigoration, a primary symptom of PD.

## Materials and Methods

### Subjects

23 PD patients [mean age 67.3 years; 12 female] and 21 matched, neurologically intact controls [mean age 61.2 years; 11 female] participated in this study. Subjects were paid according to performance on a 24-minute virtual patch-foraging, in addition to a guaranteed $10 for participation in the experiment. All subjects gave informed consent in accordance with the Institutional Review Board of the New York University School of Medicine. Patients diagnosed with idiopathic PD were referred to this study by neurologists at the New York University Parkinson’s and Movement Disorders Center (NYUPMDC). In order to be eligible for the study, patients were: determined by their neurologist to have a modified Hoehn and Yahr (HY; Hoehn and Yahr, 1967; Goetz et al., 2004) scale of motor function rating no greater than three (mild to moderate stages of disease severity); native English speakers; and on L-Dopa or DA agonists for at least three months. Patients were excluded based on a pre-test screening if they had: any history of comorbid psychiatric or neurological disturbance; any history of severe drug or alcohol abuse; a diagnosis of depression or anxiety disorder; or were taking any medication that could affect cognition other than dopaminergic medication prescribed to treat PD (Rutledge et al., 2009).

All subjects participated in two morning test sessions, approximately one week apart. The patient visits were comprised of one session on dopaminergic medication and one off. In the on session, patients were tested at least 1.5 hours after taking their medication. For the off session, they abstained from all dopaminergic medication for a minimum of 8 hours before testing. The order of on and off visits was randomized across patients.

A small number of subjects were very noisy in their responses on the task and were excluded from the subsequent analyses (Constantino and Daw, 2015). Response variance was computed within-block (over our primary dependent measure, the tree-by-tree exit thresholds, defined below) and then averaged across blocks for each visit to obtain a mean within-block variance per subject. This measure represents variation from the subject’s mean strategy and is indicative of very noisy or close to random behavior in the task. We excluded three patients and one control with response variances that fell more than 2 standard deviations above the group mean. Thus, all results reported here concern 20 patients and 20 controls (see Table 1). Of these subjects, one patient ended the task two blocks early in the off-medication condition due to physical discomfort and one control ended the second visit early due to an unexpected time constraint. Thus, these subjects are necessarily omitted from tests that turn on the last block of the affected visit but data from the unaffected visit are included where possible.

**Table 1:** Descriptive statistics for patients and controls

|  | Age | MoCA | Yrs Edu | Male (N) | HY: 0-1 | 1-2 | 2-3 | UPDRS | WASI | Yrs Onset |
| --- | --- | --- | --- | --- | --- | --- | --- | --- | --- | --- |
| control | 60.5<br>(11.6) | 27.5<br>(2.4) | 17.1<br>(2.5) | 9<br>- | - | - | - | - | - | - |
| PD | 66.2<br>(7.1) | 27.4<br>(2.1) | 17.6<br>(1.8) | 10<br>- | 4<br>- | 8<br>- | 6<br>- | 19.9<br>(10.7) | 121.2<br>(9.5) | 7.3<br>(6.3) |

HY - Hoehn and Yahr scale, counts (N); UPDRS - Unified Parkinson's Disease Rating Scale; WASI - Weschler Abbreviated Scale of Intelligence; MoCA - Montreal Cognitive Assessment. Statistics are based on $N = 20$ healthy controls and $N = 20$ patients, except for: MoCA ( $N = 16$ controls); Education ( $N = 15$ controls); HY ( $N = 18$ patients); UPDRS ( $N = 16$ patients).

**Table 2:** Parameter values defining the two environment types.

| block | Environment type |  |
| --- | --- | --- |
|  | Long travel delay | Short travel delay |
| $h$ (sec) | 4 | 4 |
| $d$ (sec) | 12 | 4 |
| $\kappa, \sigma_\kappa$ | .899, 0.09 | .899, 0.09 |
| $S_0, \sigma_s$ | 10, 1 | 10, 1 |

PD patients and matched controls did not differ significantly on the Montreal Cognitive Assesment (MoCA, a measure of mild cognitive impairment; Nasreddine et al., 2005), years of education, or age. (All p’s > 0.05; see Table 1 for details and notes on incomplete measures.) Patients were at the mild to moderate stages of disease severity, with a mean HY rating of 2^1^ (±0.8 sd) and a mean score of 19.9 (±10.7 sd) on the motor exam (section III) of the Unified Parkinson’s Disease Rating Scale (UPDRS-III; Lang and Fahn, 1989), a measure of symptom severity administered near the time of testing. The mean number of years since disease onset was 7.3 (range: 1 to 28 years). On average, patients scored 121.2 (±9.5 sd) on the Weschler Abbreviated Scale of Intelligence (WASI; Wechsler, 1999), indicating higher than population average intelligence. Disease severity outcomes are listed in Table 1.

Patients were prescribed various combinations and dosages of dopaminergic medications. Almost all subjects were receiving treatment with L-Dopa (*n* = 17), a DA precursor, and a subset of these were also taking D2 receptor agonists (*n* = 5). In addition to L-Dopa and D2 agonists, which specifically target the DA system, some patients were also taking monoamine oxidase inhibitors (*n* = 10; MAOI) and NMDA receptor antagonists (*n* = 5).

## Experiment Design and Task

Subjects made serial stay/switch decisions in a virtual patch-foraging task (Constantino and Daw, 2015): A discrete-trial adaptation of a class of tasks from the ecology literature (Charnov, 1976; Stephens and Krebs, 1986; Agetsuma, 1998; Hayden et al., 2011; Wikenheiser et al., 2013). On each trial, they were presented with a tree and had to decide whether to harvest it for apples or go to a new tree. Subjects indicated their choice by one of two key presses when prompted by a response cue. If they decided to harvest the tree, they incurred a short harvest time delay. During this time, the tree shook and the number of apples harvested was displayed, followed by a response cue prompting the next decision. As subjects continued to harvest at the same tree, the apples returned were depleted according to a randomly drawn multiplicative factor, such that on average the obtained rewards decayed exponentially.

If the subject chose to go to a new, replenished tree, they incurred a travel time delay. During this time, the old tree faded and moved off the screen and was replaced by a new tree, followed by a response cue. Each new tree had never been harvested and its initial payoff (as the starting point for the exponential decay) revealed the tree’s quality. The total time in the game was fixed. The reaction time of each choice was counted toward the ensuing harvest or travel delay so that the total interval between response cues (and thus the average reward rate) was unaffected by the response speed. Subjects who responded too slowly were penalized by a timeout lasting the length of a single harvest trial. Thus, subjects visited a different number of trees depending on their harvest decisions but were able to influence the reward rate only through their harvest or leave choices, not their reaction times.

Subjects experienced two distinct foraging environments in an ABAB counterbalanced block design. The decision-relevant parameters that define an environment are the harvest time, the travel time, the rate at which apples are depleted, and the tree quality distribution. By varying travel time across blocks, we produced two environments that differed in terms of richness or achievable average reward rate. The environment changed every 6 minutes, and this was signaled by a change in background color and a short message; the changes in the environment characteristics were not explicitly cued. Subjects were instructed that the experiment would last approximately 30 minutes, including a practice session, that trees could never be revisited, that new trees had never been harvested and were a priori identical, and that harvesting a tree would tend to return fewer apples over time. They were told that they would be paid approximately 1 cent per apple collected and should try to collect as many apples as possible.

The fixed time per block and the different time costs associated with the two actions meant that subjects had to consider both the expected rewards *and* the real time costs of each choice. Since our hypotheses concern subjects’ asymptotic choice preferences in each environment, and because examination of the response variance indicated that behavioral preferences continued to adjust through the first environment transition (and that learning about the mechanics of the experiment continued after the practice trials), we analyze only the second, more stable, occurrence of each environment type.

All results are qualitatively and directionally robust to our exclusions of both subjects and blocks. In particular, the effect of disease remains significant with all subjects and blocks included, and the medication effects are estimated in the same direction, although the standard errors increase and the effect loses significance as the high-variance initial blocks are reintroduced.

### Experiment Parameters

Each foraging environment is defined by the average initial tree richness *S*_0_, the average depletion rate per harvest *κ*, the travel time *d* and the harvest time *h*. We denote the state (current expected harvest) of a tree at trial *i* as *s_i_*. Each travel decision led to a newly drawn tree of variable quality: *s_i_* ~ 𝓝(*S*_o_,*σ_s_*) after delay *d*. Each harvest decision depleted the apples by a stochastically drawn, multiplicative decay *κ_i_*, such that *s_i+i_* = *κ_i_s_i_*, where *κ_i_* is drawn from a Beta distribution with mean κ and standard deviation *σ_κ_*. This resulted in an effective distribution of trees of varying quality with different possible reward paths. The reward earned from each harvest was a noiseless reflection of the state of the tree, *r_i_* = *s_i_*. We varied travel time *d* across blocks to create high and low average reward rate foraging environments.

### Behavioral Task Training

After reading the instructions, subjects were prompted to ask questions before beginning a 2-minute training session, in which they experienced two environments of different qualities (selected to also differ from the experimental environments). Subjects were asked to identify the shorter travel delay (richer) environment. The training task was not long enough for the patients to fully acclimate to the timing contingencies of the task prior to beginning the experiment, which, in addition to variability due to learning about the environments, likely contributed to the high response variance in the initial two experimental blocks.

### The Marginal Value Theorem and Optimal Behavior

Charnov (1976) showed that the long run reward-rate optimizing policy for this class of tasks is given by a simple threshold rule. In the context of our discrete-trial version of the task, the optimal policy is to exit a tree when the expected reward from one more harvest, *κs_i_*, drops below the opportunity cost of the time spent harvesting it. The opportunity cost of harvesting is the time spent harvesting, *h*, times the long-run average reward rate, *ρ*. The state *s_i_* of a tree is observable and equal to the received reward (*r_i_* = *s_i_*). Thus the optimal leaving rule is to search for a new tree whenever 𝔼_*s*_*i*+1__= [*r*_*i*+1_] = *κs_i_* < *ρh*.

## Threshold Estimates

Our primary dependent variable of interest was the threshold at which subjects left each tree. We estimated the thresholds as the average of the last two rewards (*r_i_* and *r*_*i*-1_) received before an exit decision. These rewards represent, respectively, a lower and upper bound on the (continuously valued) threshold since exiting at *i* implies that *κr_i_* was lower than the subject’s threshold and not exiting in the preceding decision implies that *κr*_*i*-1_ was greater. Tree-by-tree leaving thresholds are shown for two example subjects, one control and one PD patient, in Figure 2. In order to see if subjects were sensitive to manipulations of the opportunity cost of time, we compared the estimated exit threshold across the two block types, using paired t-tests on the mean (across trees, within block) per-subject leaving thresholds.

**Figure 1:**
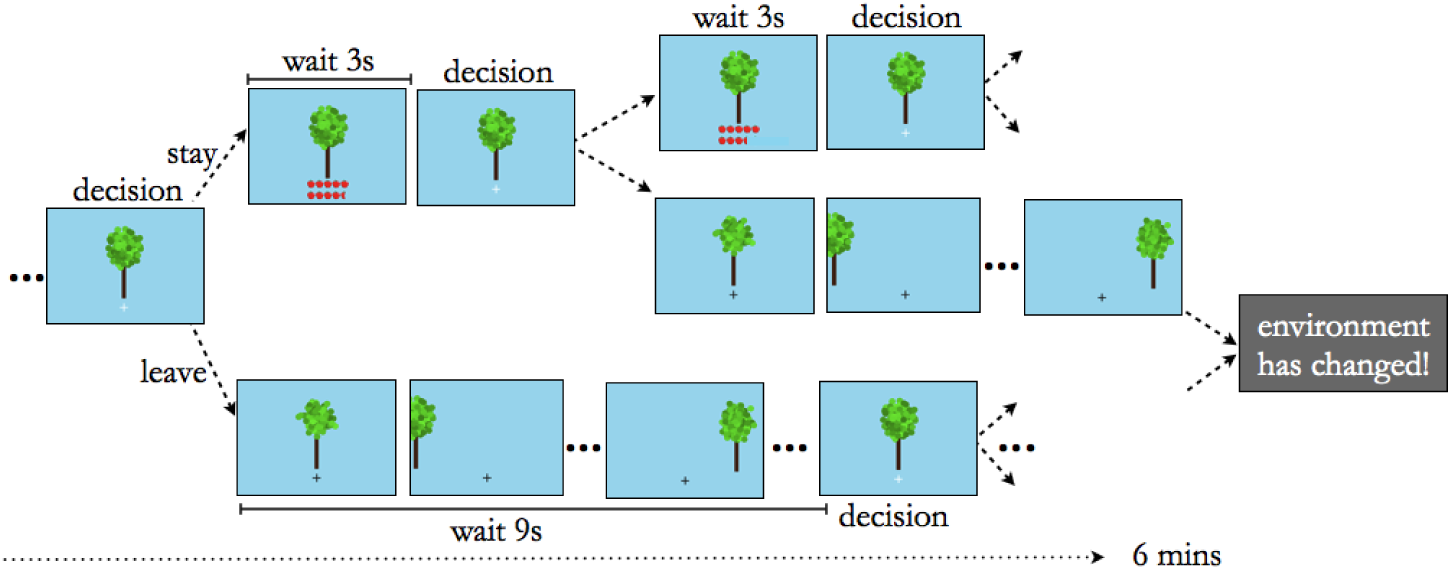
Foraging task. Subjects foraged for apples in four 6-minute virtual patch-foraging environments. They were presented with a tree and had to decide whether to harvest it for apples and incur a short harvest delay or move to a new tree and incur a longer travel delay. Harvests at a tree earned apples, albeit at an exponentially decelerating rate. New trees were drawn from a Gaussian distribution. Environmental richness (opportunity cost of time) was varied across blocks by changing the travel time. The quality of the tree, depletion rate and richness of the environment were a priori unknown to the subject (see *Methods* for a detailed explanation).

**Figure 2:**
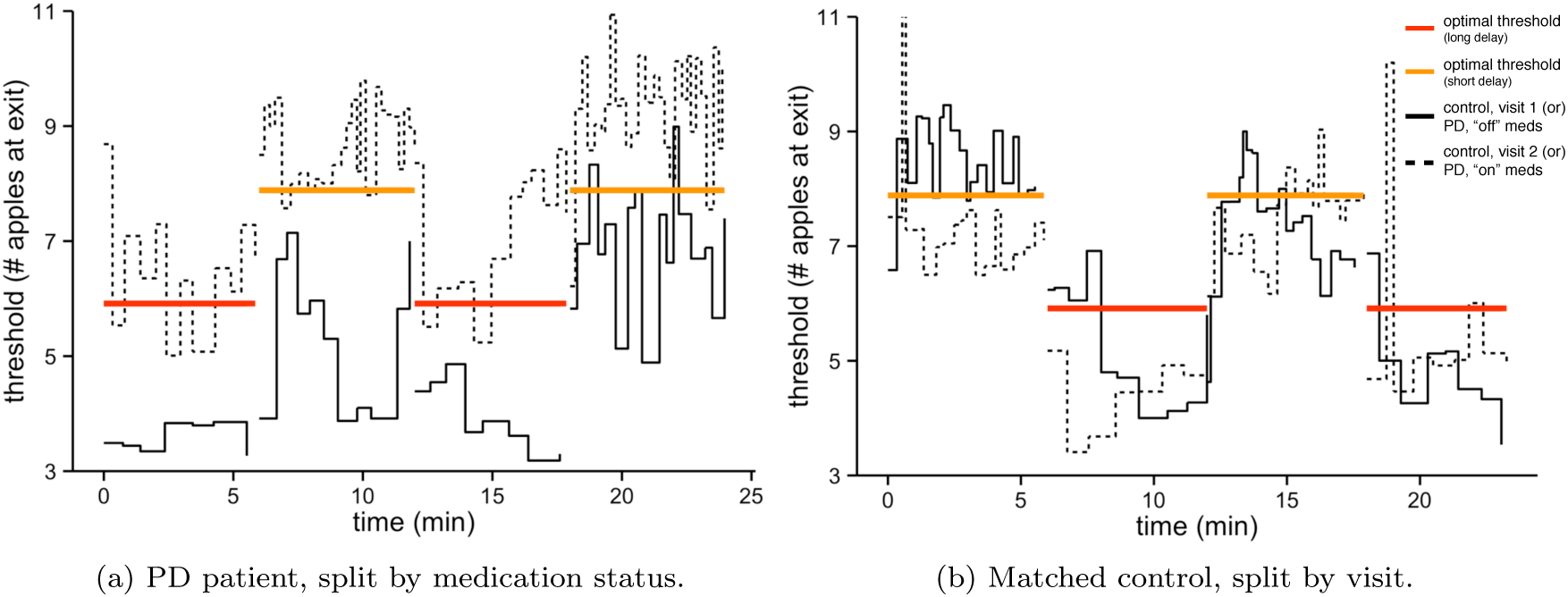
Foraging behavior – individual subject data. Example subject tree-by-tree exit points over time in the experiment (in black). Exit thresholds are in units of apples and computed as the mean number of apples eraned in the last two harvests before an exit decision (see Methods). The dotted line indicates the on medication thresholds for the patient (a) and the second visit thresholds for the control (b). Colored lines are the optimal exit threshold in the short (orange) and long (red) travel delay environments.

## Effect of Disease and Medication on Leaving Thresholds

We also examined how thresholds varied with disease and medication status. In order to test this across subjects, while controlling for both between- and within-subject variation and the repeated measures structure of the task, we used a linear mixed effects model, estimated using the lme4 package in the R statistical language (Bates et al., 2011). We computed p-values for coefficient estimates using the lmertest package, which uses the Satterthwaite approximation to estimate the degrees of freedom (Kuznetsova et al., 2014). For this analysis, the dependent variable was the tree-by-tree exit thresholds and the explanatory variables were binary indicators for visit (to control for test-retest effects), environment type (long or short), disease, medication status, and the interactions of the disease and medication effects with block. The within-subject factors — intercept, visit, environment-type, medication, and the interaction of environment with medication — were included as random effects (i.e. allowed to vary across subjects). Disease (0: patient; 1: control) and medication (0: patient off and controls; 1: patient on) were coded such that the baseline captures patients off medication and the disease and medication coefficients capture the two main comparisons of interest, that between controls and unmedicated patients (the between-subject effect of disease, off medication) and that within patients on vs. off medication (the within-subject medication effect in patients), respectively.

As a robustness check, we ran variants of this regression that included nuisance variables, such as age, education and MoCA.

## Control Task for Motor Perseveration

In order to test for potential motor perseveration, on each visit we included a control task after the foraging task that lasted the length of one foraging block (Figure 4a). 19 subjects completed this task: 9 controls [mean age 61.22 years; 4 females] and 10 patients [mean age 66.2 years; 3 females]. Due to late introduction of the perseveration control, this sample only partially overlapped the participants in the foraging task; specifically, 6 controls and 5 patients were tested alongside the foraging task in the same visits, whereas the remaining subjects were newly recruited (from the same referral population) as part of a different study. This sample of patients had a mean HY rating of 2.11 (±0.65), a mean UPDRS-III of 24.11 (±9.9), and a mean WASI of 117.1 (±10.61).

The perseveration task had the same timing and response latencies as the harvest decisions in the foraging task but there were no rewards and no manipulation of the opportunity cost of time. On each trial, subjects saw a randomly drawn shape (diamond or star; drawn with equal probability) displayed in the center of the screen and were prompted to indicate the shape by a shape-specific key press (see Figure 4a). The shape was displayed for the same amount of time as the apples in the foraging task. A failure to respond resulted in a timeout, and all responses were followed by feedback (“correct”, “incorrect”, or “too slow”). Perseveration in motor responses would appear here as a tendency to press the same key as in the preceeding trial, independently of the presented shape in the current trial.

We investigated whether response accuracy was affected by motor perseveration using a mixed effects logistic regression. The dependent variable was a per-trial indicator of whether the response was correct, and the explanatory variables were indicators for whether the shape had changed (relative to the previous trial) and for visit. All variables, including the intercept, were taken as random effects across subjects. We repeated the regression, this time replacing the dependent variable with an indicator for trials in which the subject timed out. Finally, we also tested for an overall effect of disease and medication on correct responses by including those indicators, as well as one for visit, as dependent variables.

## Results

PD patients and healthy controls completed a computerized decision task in which they repeatedly chose whether to harvest apples from a tree whose return gradually depleted, or seek a new, replenished tree. On each visit, subjects faced two types of foraging environments, differing in richness, in alternating order across four blocks. Environmental richness was decreased by increasing the travel delay between trees, which, all else held equal, reduced the rate at which apples could be earned. This reduction in potential earnings decreased the opportunity cost of time spent harvesting. An ideal forager should harvest each tree more extensively as the opportunity cost of time decreases, even if it means harvesting for smaller rewards (down to a lower earned apple threshold), because time is relatively cheap and rewards are relatively scarce.

The literature linking the average reward rate, a measure of the opportunity cost of time in our task, to the DA system, suggests that lower tonic DA levels would result in a lower estimate of environment quality, all else held equal. In our study, this hypothesis would imply that PD patients off medication should harvest trees down to lower leaving thresholds, whereas medication should increase thresholds.

Individual leaving thresholds over the course of the experiment are shown for two example subjects, one control and one PD patient, split by visit and medication status (on/off), respectively (Figure 2). Subjects successfully entered an average of 341 harvest-or-exit decisions at an average of 50 trees per session, and failed to indicate a decision within the allotted time on an average of 5 trials (1.5% of trials). The frequency of a timeout did not differ significantly between control and patient groups (*t*_38_ = −1.12, *p* = 0.269), even though reaction times were significantly slower in the patient group (*t*_38_ = −2.688, *p* = .011). Reaction times within the patient sample did not differ with medication status. Importantly, events in the task were scheduled such that faster or slower reaction times would not affect the timing of any subsequent task events and rewards (and therefore could not affect earnings or the optimal choice policy), so long as responses occurred within the allotted time window.

PD patients earned similar amounts on ($17.7) and off ($17.0) dopaminergic medication but on average earned significantly less than matched controls ($18.7; *t*_38_ = 2.614, *p* = 0.013). Finally, controls did not differ significantly on any measure across the two visits, other than in the mean number of late-response warnings and mean reaction time, which were both significantly lower in the second visit (warnings: *t*_19_ = 2.21, *p* = .04; rt: *t*_19_ = 3.11, *p* = .006).

## Foraging Behavior: Compliance With the MVT

The initial quality of each new tree and the depletion rate following a harvest decision were randomly drawn. These features require the ideal agent to monitor the number of apples obtained at each step, which serves as a noiseless measure of the current state of the tree, in order to decide whether the current tree is worth harvesting. The optimal policy for deciding whether to leave a tree requires comparing the expected reward from the next harvest to a threshold, which is controlled by the overall average reward per timestep, i.e. the opportunity cost of time in the environment. Subjects should harvest trees more thoroughly (down to lower returns) when the overall reward environment is poorer; in this sense the observed level of harvesting reflects an evaluation about the quality of the environment.

Thus, we examined subjects’ exit thresholds — estimated from the number of apples received on the last harvests before leaving a tree — as our primary dependent measure. The threshold data of an example control and patient (Figure 2) show gross threshold adjustments across environments in the direction predicted by the optimal analysis: They decrease their leaving thresholds in the lower quality environment, reflecting the lower opportunity cost of time. This pattern is consistent with the average behavior within each group (Figure 3). Paired t-tests find that the leaving threshold adjustments were significant and in the predicted direction for the control group (visit 1: *t*_19_ = −7.72, *p* < .001; visit 2: *t*_18_ = −6.4, *p* < .001) and patient group, both on medication (*t*_19_ = −4.88, *p* < .001) and off (*t*_18_ = −3.01, *p*= .008)^2^.

**Figure 3:**
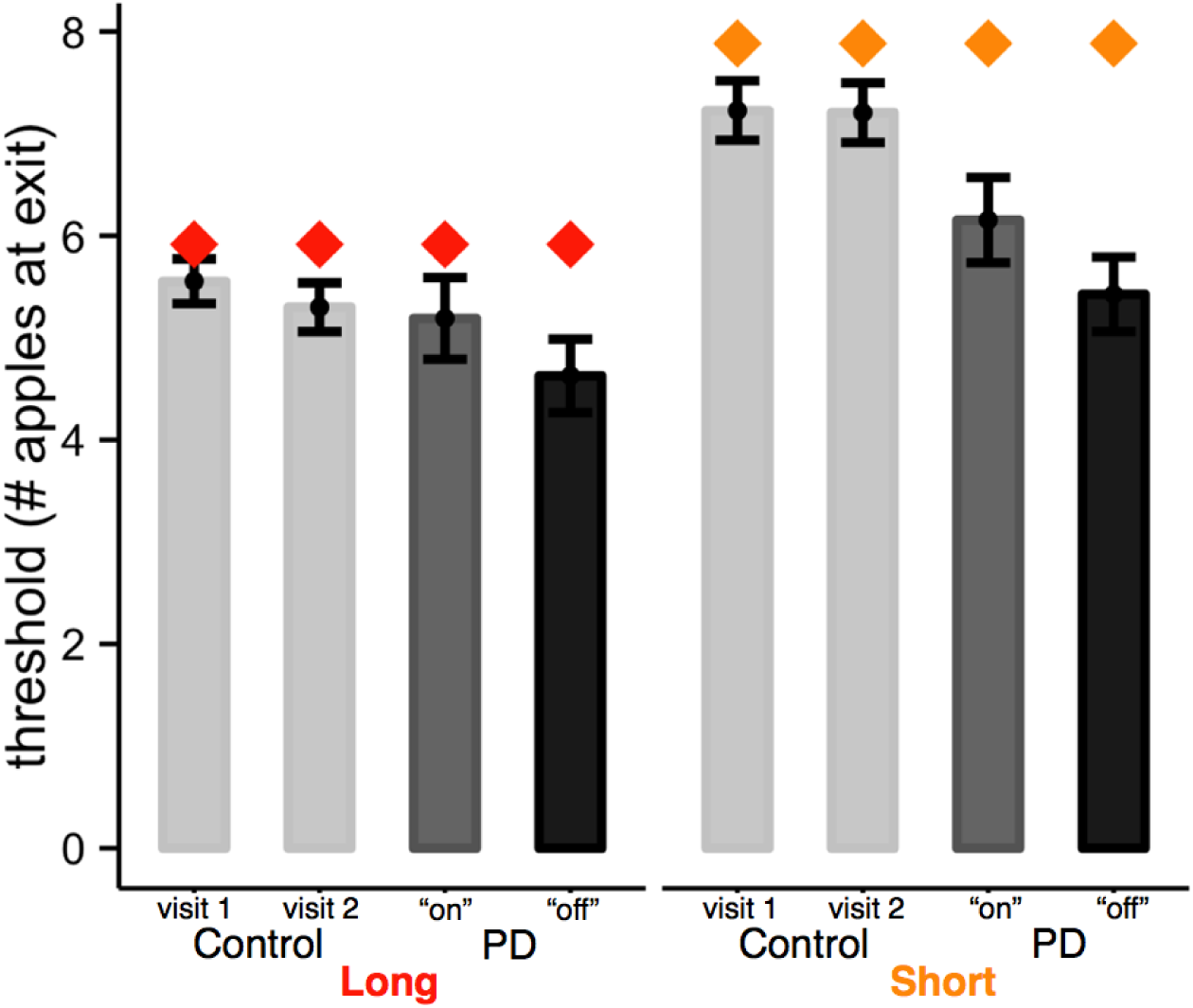
Foraging behavior – group data. Group exit thresholds by environment type, visit and medication status. The grey bars indicate mean leaving thresholds per environment type (long or short), with 95% confidence intervals, displayed in the following order (left to right): controls, visit 1; controls, visit 2; PD patients, on medication; PD patients, off medication. Filled diamonds represent the ideal forager thresholds for short (orange) and long (red) travel delays.

Comparing the numerical thresholds (averaged across visits) to the specific values predicted under optimal switching, we found that both groups had a tendency to harvest longer (i.e. exhibit a lower exit threshold) than optimal in all environment types (control, long: *t*_19_ = −2.134, *p* = .046; control, short: *t*_19_ = −2.23, *p* = .038; disease, long: *t*_19_ = −2.55, *p* = .019; disease, short: *t*_19_ = −5.66, *p* < .001), which resonates with the findings of a previous study examining the behavior of healthy subjects in this task (Constantino and Daw, 2015). Altogether, these results suggest that subjects adust their behavior to changes in the opportunity cost of time in the direction predicted by theory, though with a bias to over-stay relative to the optimal policy.

## Foraging Behavior: Effects of Disease and Medication

All subject groups, irrespective of disease, visit or medication condition, adjusted their leaving thresholds to changes in the opportunity cost of time occasioned by manipulations of environmental quality. In order to examine any between-group differences in these adjustments, while accounting for within and between subject variance, we analyzed subjects’ tree-by-tree leaving thresholds using a linear mixed-effects regression. This analysis indicated that PD patients, when off medication, harvested significantly longer (i.e., displayed lower exit thresholds) than controls (*t*_36_ = 2.5, *p* = .017). Medication significantly ameliorated the deficit: Patients had higher exit thresholds on medication as compared to off (*t*_16_ = 3.09, *p* = .007) (see Table 3).

**Table 3:**
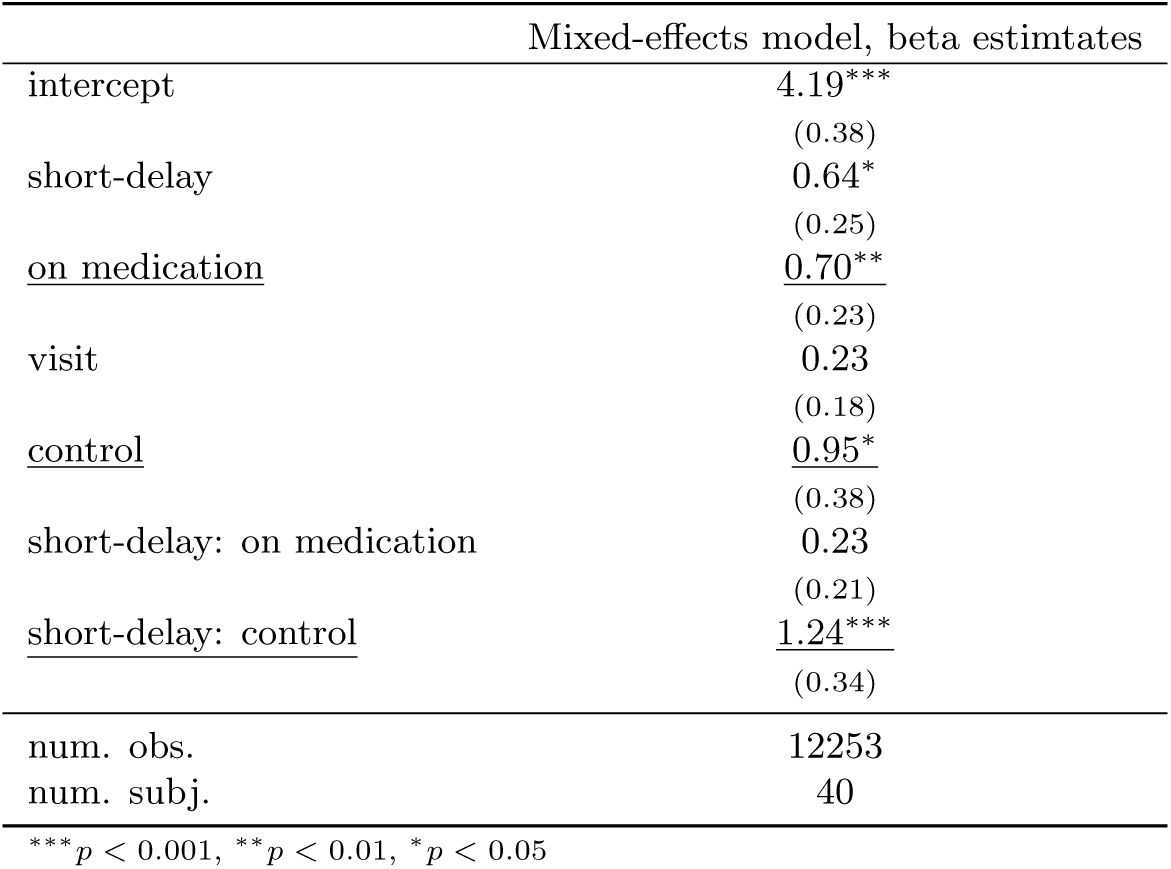
Effects of disease and medication on foraging behavior

Consistent with the hypothesis that DA tracks the opportunity cost of time, these results suggest that lower DA levels in the unmedicated patients resulted in longer harvesting — as though the estimated opportunity cost of harvesting was biased downward — and that DA replacement therapy remediated this effect. Note that although the pharmacokinetics are clearly complex, if we envision disease and medication as each exerting something like a multiplicative gain on the tonic DA level (e.g., changing the constant of proportionality by which, hypothetically, it reflects the average reward rate), then we would expect these manipulations to exert both the overall effects discussed thus far and also an interaction with block type (a proportionally more prounounced decrease in richer environments) (Beierholm et al., 2013). Indeed, the effects of both drug and disease were most apparent in the richer (short travel-delay) environment, as shown in Figure 3, though this interaction was only significant for the effect of disease (*t*_34_ = 3.67.*p* < .001).

## Motor Perseveration and Other Controls

Our hypothesis – that biased signaling of the opportunity cost of time in Parkinsonism leads to a failure to disengage from low-paying options – amounts to the prediction that subjects suffer a form of cognitive perseveration on harvesting. This is different from, but might be confounded with, simpler reward-independent perseveration that could arise from gross motor difficulties – a characteristic symptom of PD – when moving between the response keys. (The association of keys with harvest or exit was, for simplicity, not randomized across trials.) We designed the task with relatively slow timing requirements in order to mitigate any movement difficulties that could otherwise trivially favor perseverative responding. To investigate whether motor perseveration contributed to foraging behavior, we examined an additional forced-choice control task with the same timing and key press requirements as in the foraging task but without the opportunity cost and reward components.

In the task, subjects responded to two randomly drawn shapes with shape-specific key presses (see Figure 4a). On each trial, a random shape was drawn from two possible alternatives, resulting in unexpected sequences of the same shape followed by a change in shape. Reward-independent motor perseveration would appear as a heightened tendency to press the key associated with the previously presented shape (a motor-response stickiness), and thus a higher error rate on trials that require a switch in the motor response (change in shape).

**Figure 4:**
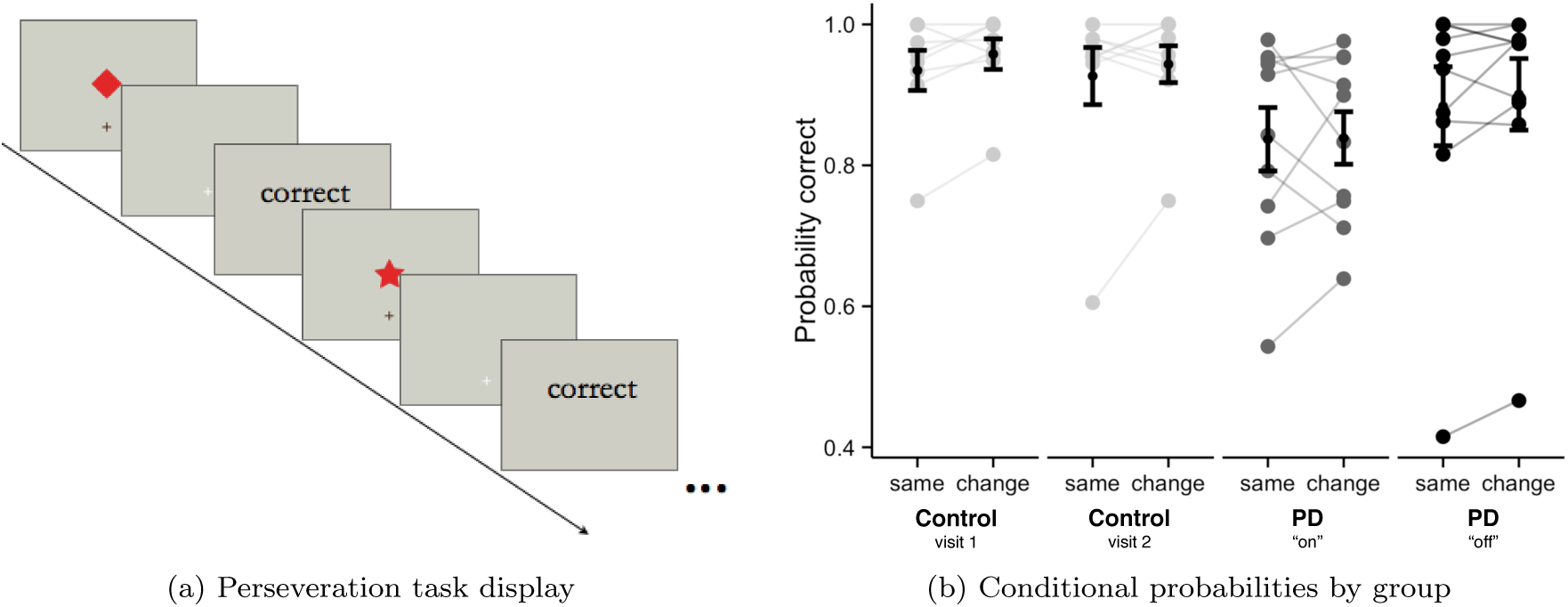
Motor perseveration task. (a) The perseveration task lasted the length of one foraging environment (6 minutes). During this time, one of two shapes (diamond or star) was randomly drawn (with equal probability) and displayed center screen on each trial, resulting in unexpected sequences of the same shape followed by a change in shape. The shape remained on screen for the same amount of time as the apples in the foraging task and was followed by a response cue and feedback (“correct”, “incorrect”, or “too slow”). The timing and keypress contingencies were the same as in the foraging task but now one of the shapes was randomly assigned to the “harvest” key and the other to the “exit” key. (b) Probability of a correct response following the consecutive presentation of the same shape and following a switch in shape, shown by group in the following order (left to right): controls, visit 1; controls, visit 2; patients, on medication; patients, off medication. Individual points connected by grey lines are individual subject probabilities. Error bars show the 95% confidence intervals around the group mean probability.

We did not find a significant effect of shape change on the probability of being correct in any of the groups (patients on medication, patients off medication, and controls; see Figure 4b and Table 4). Correct responses overall were lower for patients on medication but this main effect was independent of the shape sequence, indicating it did not arise from perseveration. Furthermore, as this deficit *increased* with medication, it ran in the opposite direction of the increased tendency for harvesting by patients off medication in our foraging task. We repeated the same logistic regression analysis, this time replacing the correct response dependent variable with an indicator identifying missed trials, and again found no effect of a change in shape for any of the groups (results not shown). Together, these negative results suggest that motor perseveration did not contribute appreciably to the effects of disease and medication on the foraging task.

**Table 4:** Motor perseveration fits

|  | PD, off meds | PD, on meds | Controls |
| --- | --- | --- | --- |
| intercept | 2.95***<br>(0.86) | 1.57***<br>(0.34) | 2.68***<br>(0.45) |
| <u>change</u> | <u>0.24</u><br>(0.25) | <u>0.00</u><br>(0.19) | <u>0.34</u><br>(0.23) |
| visit | -0.07<br>(1.11) | 0.83<br>(0.53) | 0.55*<br>(0.27) |
| num. obs. | 874 | 882 | 1482 |
| num. subj. | 10 | 10 | 9 |

\*\*\* $p < 0.001$ , \*\* $p < 0.01$ , \* $p < 0.05$ , PD: Parkinson's disease

## Discussion

PD patients and matched controls performed a virtual patch-foraging task that measured how long they stayed with a depleting resource. We hypothesized that choices would be modulated by tonic DA, which has been suggested to carry the key decision variable for this class of tasks: The average reward rate or opportunity cost of time (Niv et al., 2007; Cools et al., 2011). Consistent with DA deficiency reducing the subjective opportunity cost of time, PD patients off medication harvested down to a lower reward threshold than controls, and DA replacement medication alleviated this deficit. These results suggest a neural substrate for stay/switch decisions or stopping problems (like foraging), where options are encountered serially rather than simultaneously, and propose a novel mechanistic link between choices in this class of tasks and the continuous modulation of behavioral vigor, which has also been linked to a putative dopaminergic opportunity cost signal (Niv et al., 2005, 2007; Guitart-Masip et al., 2011; Salamone and Correa, 2012; Beierholm et al., 2013).

Drawing on parallel opportunity cost considerations, previous research has suggested that tonic DA might control behavioral vigor (Niv et al., 2005, 2007). This hypothesis helps to unify research on DA’s role mediating prediction errors and learning about choices from rewards with the neuromodulator’s more direct effects on activation and movement. Although outcomes in the present task are, by design, unaffected by vigor (reaction times), the choices themselves turn on the opportunity cost of the time intervals associated with the different actions. This design allows us to investigate the role of opportunity costs directly without relying on reaction times or assumptions about unobserved energetic costs of vigorous action (Niv et al., 2005; Guitart-Masip et al., 2011). Neurally, it has been proposed that tonic DA governs the vigor of actions directly, perhaps by modulating the direct and indirect pathways out of striatum (DeLong, 1990). The present results suggest that a common mechanism controls the more discrete decision of whether (or when) to leave an option in search of something better. This effect could be subserved by analogous dopaminergic modulation of the pathways in a different striatal subregion.

An alternative target for DA’s effects is the anterior cingulate cortex, which other studies have shown to play a key role in foraging decisions (Hayden et al., 2011; Kolling et al., 2012; Rushworth et al., 2012; Mobbs et al., 2013; Blanchard and Hayden, 2014) (but see Shenhav et al., 2014). Suggestively, the cingulate has also been associated with cognitive control, and recent research has proposed that this seemingly disjoint function is itself governed by opportunity cost considerations about how to allocate limited neural resources (Keramati et al., 2011; Kurzban et al., 2013; Shenhav et al., 2013; Boureau et al., 2015).

One important question concerns how our result relates to previously observed reinforcemement learning deficits in PD. Many earlier studies have demonstrated deficits using choice tasks that require learning about the values of different competing actions (Cools et al., 2001; Frank et al., 2004; Shohamy et al., 2004; Bódi et al., 2009). In order to minimize the role of learning, we focused on asymptotic behavior in the second block of a stable task. That said, although our hypothesis focuses on tonic DA, neither PD nor its medications specifically target tonic rather than phasic DA (but may plausibly affect both), and indeed many previous results have been interpreted in terms of phasic DA’s widely hypothesized role in signaling prediction errors for action value learning (Frank et al., 2004; Rutledge et al., 2009). In particular, it has been suggested that deficits in phasic DA signaling emphasize avoidance learning or learning from negative prediction errors, with medication, conversely, favoring rewards (Frank et al., 2004; Rutledge et al., 2009). However, the application of these principles to the sequential foraging problem does not offer a straightforward prediction.

A key qualitative difference is that our task does not directly pose an action-value learning problem, since options are encountered one at a time and the decision is when to reject an option in favor of searching for a better one. Our task can be reframed in the simultaneous choice terms of those other accounts by focusing on the learning of the long-run expected values for the two competing responses (harvesting vs. searching for a new tree), though we have previously shown that this framework does a worse job of explaining behavior than choices based on tracking only the opportunity cost of time (Constantino and Daw, 2015). In any case, framed this way, a bias toward learning from negative prediction errors should depress both options and would not appear to predict the observed asymmetric behavioral bias toward harvesting longer.

Alternatively, our hypothesis, informed by related work on action vigor (Niv et al., 2005) and foraging (Kolling et al., 2012), is that choices in this class of problems are categorically different from simultaneous choice in that they are governed by a single global decision variable: The overall average reward rate, which is the opportunity cost of engaging the current option. This opens up the possibility that performance is mediated by a more global neural variable, such as tonic extracellular DA (Niv et al., 2005; Beierholm et al., 2013). Here and in the vigor research, (tonic) DA is hypothesized to be involved directly in the <u>expression</u> of the behavior, rather than only indirectly, via controlling plasticity. Although our data do not directly speak to this point, results from a study manipulating drug status independently during both training and test in a reward learning task suggest that apparent PD learning deficits are actually driven by expression rather than learning, which supports the present framework and is hard to understand given the standard interpretation involving deficits in phasic prediction error signaling (Shiner et al., 2012).

As in any PD study, our patient and control populations were somewhat heterogeneous and within the patient population there was a range in the combinations and dosages of dopaminergic medication. These differences are unlikely to confound our results since demographic characteristics were not significantly different between patient and control groups, and the on/off medication comparison was within-subject. Furthermore, as a robustness check, we repeated the key regression but included different additional factors as nuisance variables (e.g. age, education, MoCA; data not shown) and found that the results were unaffected.

In addition to the average reward, there are several other possible mechanisms that might in principle mediate the effects we observed. In particular, DA-dependent effects on time discounting or risk preferences might also affect willingness to incur a delay in seeking a new tree of unknown value. However, the relationship between DA and those preferences is not especially consistent (see Cools et al. (2011) for review). Moreover, since the average reward can affect preferences in both intertemporal choice (via the opportunity cost of time) and risky choice (via reference points and framing effects), as well as vigor and perseveration, the opportunity cost hypothesis may provide a more general account of DA function (Cools et al., 2011). However, it remains for future work to independently assess risk sensitivity, time preferences, and foraging behavior, in order to understand how they relate to one another and to DA dysfunction.

Similar points apply to interpreting our results in terms of perseveration. Although we took steps in both task design and a control experiment to rule out an explanation in terms of pure motor perseveration, the observed tendency of patients to stay rather than switch trees is reminiscent of cognitive perseveration and switching deficits that have been well documented in PD patients (e.g. in task set or rule switching) (Cools et al., 2003). The average reward mechanism suggested here proposes a quantitative, underlying explanation for such dopaminergic related changes in switching behaviors, both in our serial decision task and more generally: As DA levels are depleted, they signal impoverished environments, which in turn make any status quo option appear relatively better (Cools et al., 2011). While it is difficult to map serial foraging decisions directly onto explicit tests of cognitive control and switching, there is a parallel between the cognitive perseveration (decreased flexibility or a tendency to stick to the same strategy; for example, as measured by the Wisconsin Card Sorting Task; Heaton et al., 1993) observed in PD and the increased tendency to harvest (or decreased propensity to switch) observed in our foraging task.

Indeed, the present results may ultimately point toward a similar account of internal, cognitive switching decisions of this sort and of cognitive control phenomena more generally. This would add to, and hint at a potential neural mechanism for, an emerging understanding that decisions about the allocation of one’s cognitive resources involve rational cost/benefit tradeoffs that are in many ways analogous to evaluating foraging opportunities (Keramati et al., 2011; Kurzban et al., 2013; Shenhav et al., 2013; Boureau et al., 2015).

## Acknowledgments

This research was funded by Human Frontiers Science Program Grant RGP0036/2009-C and grant R01MH087882 from the National Institute of Mental Health. Nathaniel Daw is supported by a Scholar Award from the McDonnell Foundation. We would like to thank Elizabeth A. Phelps, Jian Li and Peter Sokol-Hessner for valuable insights and discussion. The authors declare no competing financial interests.

## Footnotes

1 A fraction of patients were referred by external neurologists in accordance with our criteria. For two of these individuals we only know that the HY rating was within the 0 to 3 range; For four of these individuals we were unable to obtain the UPDRS scores.

2 The missing degree of freedom for visit 2 and the off medication condition is due to early termination of the experiment in one of the visits by one control and one patient. See methods for details.

